# The fetal hydrops-associated single-residue mutation L322P abolishes mechanical but not chemical activation of the PIEZO1 ion channel

**DOI:** 10.1101/2025.02.18.638930

**Authors:** Jinghui Jiang, Wei Guo, Xudong Chen, Yubo Wang, Xiaohui Zhu, Liying Yan, Jie Qiao, Bailong Xiao

## Abstract

The mechanically activated PIEZO1 ion channel is genetically linked to numerous physiological and pathophysiological processes. For example, deleting *PIEZO1* in mice leads to defective lymphatic vessel development, while nonsense mutations in humans are associated with autosomal recessive generalized lymphatic dysplasia (GLD) and non-immune hydrops fetalis. However, it remains unclear whether PIEZO1-dependent biological processes are directly mediated by its intrinsic mechanosensitivity. Here we identified a human fetal hydrops-associated single-residue mutation, L322P (corresponding to L329P in mouse PIEZO1), which completely abolishes PIEZO1’s response to mechanical stimuli while maintaining normal plasma membrane expression and responsiveness to its chemical activators, Yoda1 and Jedi1. Remarkably, the mechanical response of the mutant can be restored by Yoda1. These findings demonstrate a direct link between the loss of PIEZO1’s mechanosensitivity and the pathophysiological phenotype of fetal hydrops, and raise the therapeutic potential of using PIEZO1 chemical activators to restore the mechanosensitivity of PIEZO1 missense mutants that are associated with genetic diseases such as GLD and hydrops fetalis.

**Significance Statement:** Genetic deletion studies have revealed the diverse cellular, physiological, and pathophysiological roles of the mechanically activated PIEZO1 ion channel. However, whether these functions are directly mediated by its mechanosensitivity has remained experimentally unaddressed. In this study, we identified a novel fetal hydrops-associated single-residue mutation, hPIEZO1-L322P, which specifically abolishes PIEZO1’s mechanical response while retaining its chemical activation. Remarkably, the mechanical response of the mutant can be fully restored by the PIEZO1 chemical activator Yoda1. These findings address the long-standing question of whether PIEZO1 relies on its mechanosensitivity to mediate in vivo functions, provide critical insights into the mechanosensing mechanism of PIEZO1, and highlight the therapeutic potential of using PIEZO1 chemical activators to rescue the loss of mechanosensitivity caused by missense mutations.

## Introduction

PIEZO1 functions as a versatile mechanotransduction channel for converting different forms of mechanical stimuli such as membrane indentation, stretch and shear stress into Ca^2+^ signaling in various cell types such as endothelial cells, red blood cells, cardiomyocytes, and astrocytes ^1–10^. Genetic deletion of *PIEZO1* in specific cell types in mice have demonstrated its wide variety of cellular, physiological and pathophysiological processes involving mechanotransduction ^7,11^. For instance, lymphatic endothelial cell-specific deletion of *PIEZO1* leads to defective development of the lymphatic vessel system ^12,13^. Importantly, many of the PIEZO1-mediated physiological function revealed from animal models have been verified in human genetic diseases derived from *PIEZO1* mutations ^14–19^. Loss of the full-length PIEZO1 protein due to either nonsense or splicing site mutations in human *PIEZO1* has been linked to autosomal recessive generalized lymphatic dysplasia (GLD) with non-immune hydrops fetalis ^19,20^. Thus, the cellular, physiological and pathophysiological roles of PIEZO1 have been strongly supported by both mouse and human genetics studies.

PIEZO1 is a large membrane protein with 38 transmembrane helices (TM) ^7,21,22^. For instance, human and mouse PIEZO1 (hPIEZO1 and mPIEZO1) have 2522 and 2547 residues, respectively ^1^. Cryo-EM structural studies have revealed the modular organization of the complex 38-TM-topological structure of PIEZO1 ^23–26^. The N-terminal 36 TMs are folded into 9 repetitive transmembrane helical units (THUs) and form the highly curved blade domain, which is proposed to function as the mechanosensing domain of PIEZO1 ^23,27^. The rest of the C-terminal domain with the last 2 TMs (termed outer helix (OH) and inner helix (IH), respectively) trimerizes to form the ion-conducting pore module ^23,24,28^. Force-induced conformational changes of the mechanosensing blade domain are associated with the gating of the ion-conducting pore ^29,30^. Most of the GLD-causing mutations lead to truncation of the C-terminal pore domain and thus are expected to generate non-pore-containing PIEZO1 proteins ^19,20^. Several GLD-associated missense mutations of *PIEZO1* result in reduced expression in the plasma membrane, and consequently leading to loss-of-function of PIEZO1^20^. On the basis of the force-induced large deformation of the PIEZO1-membrane system ^29,31,32^, PIEZO1 might directly signal their conformational changes in addition to its ion channel function ^33^. Thus, whether PIEZO1-dependent physiological and pathophysiological responses are mediated by its mechanosensitivity remains inconclusive.

Here we report the identification and characterization of a novel fetal hydrops-associated missense mutation hPIEZO1-L322P that specifically abolishes the mechanical response, but retains chemical activation of PIEZO1, which provides compelling evidence supporting the direct effect of the mechanosensitivity of PIEZO1 in determining its physiological and pathophysiological functions.

## Results

### Identification of the hPIEZO1-L322P mutation and E997-truncation in association with fetal hydrops

One clinical case in this study involves a recurrent fetal malformation characterized by fetal hydrops and lymphatic dysplasia. This young couple had experienced recurrent abortion for three times within seven years, the fetus consistently presented with hydrops at 24 weeks of gestation, leading to Mid-trimester induced abortion. After clinic evaluation, both of them considered healthy without any clinical symptoms including blood system diseases. No relative medical conditions and similar obstetric and reproductive history in both family of this couple after interview (Figure 1A). The fetal tissue obtained from the last abortion was available for sequential genetic testing. After ruling out the possibility that fetal hydrops was caused by chromosome number variation abnormalities (Supplementary Figure 1A), we performed Trio-based whole-exome sequencing (Trio-WES) for this family (Figure 1A and Supplementary Figure 1B, C). The result showed that the fetus carried compound heterozygous variants on *PIEZO1* (NM_001142864) inherited from parents respectively: c.2991+1 G>A (paternal, according to the ACMG guidelines classified as likely pathogenic: PVS1+ PM2-supporting) and c.965 T>C (maternal, according to the ACMG guidelines classified as VUS: PM2-supporting, PP3-Moderate) (Figure 1A and Supplementary Figure 1B, C). Both identified variants have not been previously reported.

**Figure 1.**
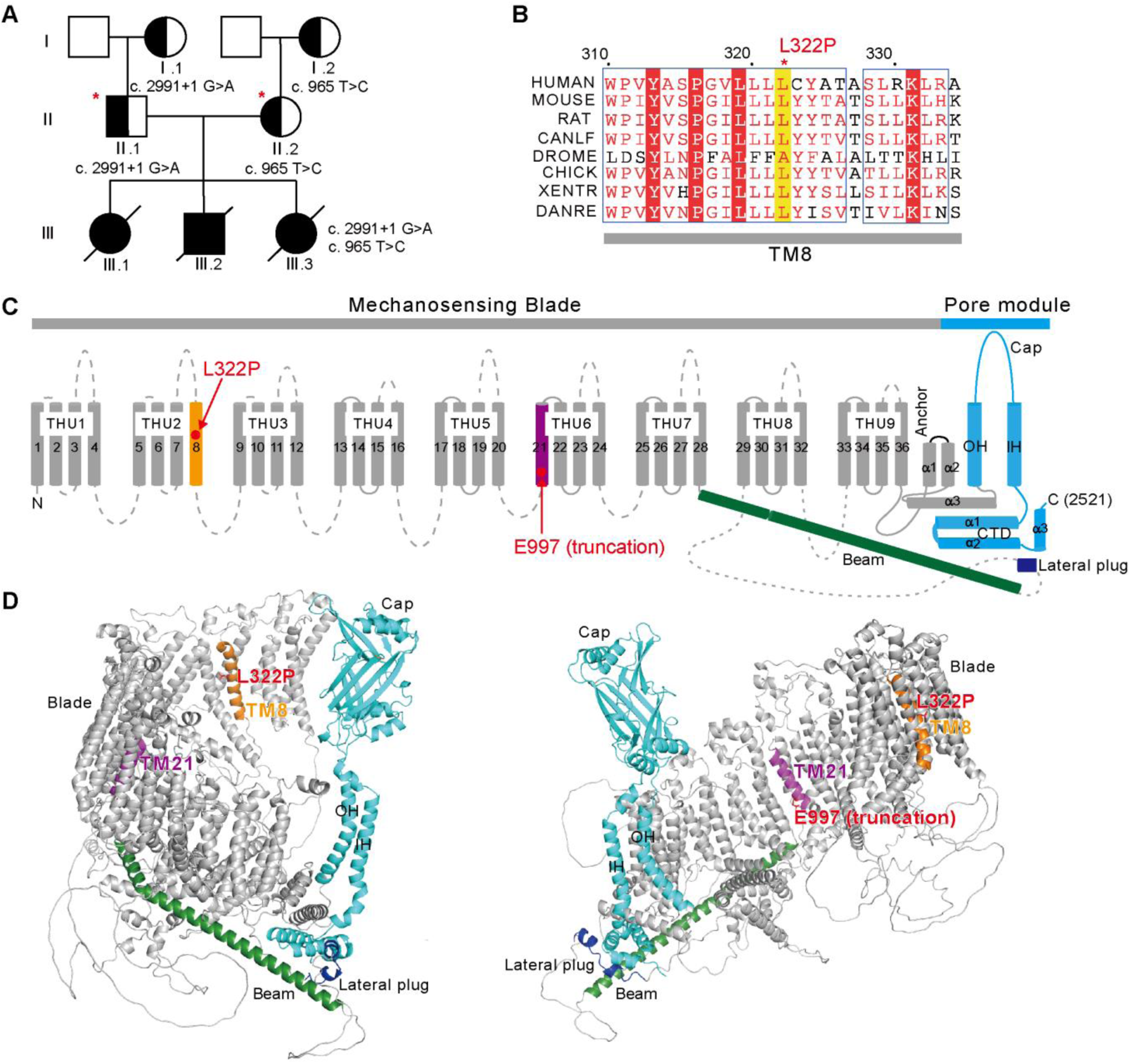
Identification of the fetal hydrops-associated hPIEZO1-E997 (truncation) and hPIEZO1-L322P mutations. **A**, Pedigree of proband family (male as circle, female as square) showing the distribution of the indicated novel hPIEZO1 variants. **B**, Protein sequence alignment of the portion containing L322 of hPIEZO1 from different species, illustrating the conservation of L322. **C**, Schematic representation of the location of the L322P and E997 truncation mutations in the 38-TM-topological structure of hPIEZO1, which include the N-terminal mechanosensing blade comprising 9 repetitive transmembrane helical units (THU1-9) and the helical beam, and the C-terminal pore module containing the outer helix (OH), Cap, inner helix (IH) and C-terminal domain (CTD). **D**, The AlphaFold2-predicted full-length structure of a hPIEZO1 protomer containing 38 TMs with the highlighted domains and the location of the L322P and E997 truncation positions. The hPIEZO1 protomer trimerizes to form the three-bladed and propeller-shaped structure.

PIEZO1 mediates mechanotransduction in lymphatic endothelial cells^12,13,34^. Importantly, previous studies have identified that loss-of-function (LOF) mutations in hPIEZO1 are associated with autosomal recessive generalized lymphatic dysplasia (GLD) with non-immune hydrops fetalis^19^. The variant c.2991+1 G>A alters the canonical splice donor motif of exon 21, and is predicted to result in the inclusion of the 424-base-pair (bp) intron 21 in the mature messenger RNA, which leads to a premature stop at the residue E997 located at the TM21 of the THU6 and truncation of the C-terminal pore domain of hPIEZO1 protein (Figure 1C, D). Thus, the variant is expected to cause a LOF of the hPIEZO1 channel.

The c.965 T>C variant induced the substitution of the highly conserved Leucine-322 residue into Proline in PIEZO1 protein (hPIEZO1-L322P) (Figure 1B). The L322 residue in hPIEZO1 and the corresponding residue L329 in mPIEZO1 are located in the TM8 of the THU2 of the N-terminal mechanosensing blade domain (Figure 1C, D). Due to that the distal THU1-3 portion of both mPIEZO1 and hPIEZO1 remain structurally unresolved, we adopt the AlphaFold2-predicted full-length structure of human PIEZO1 with the full 38-TM-topology and show the location of the L322P mutation and the expected truncation site E997 (Figure 1C, D). The blade domain has been proposed to serve as the mechanosensing domain for conferring mechanosensitivity to PIEZO channels ^23,28,35^. We hypothesized that the rare and previously unreported hPIEZO1-L322P missense mutation might abolish the mechanical activation of hPIEZO1, and consequently in combination with the LOF truncation variant of PIEZO1-E997-truncation might underlie the genetic cause of the observed fetal hydrops.

### The L322P mutation abolishes mechanical activation of hPIEZO1

To characterize the effect of the L322P mutation on hPIEZO1-mediated mechanically activated currents, we generated the hPIEZO1-L322P mutant and heterologously expressed the wild-type hPIEZO1 and the mutant in human embryonic kidney 293T cells in which the endogenous PIEZO1-encoding gene was knocked out (HEK-P1-KO cells)^36^ (Figure 2). In the whole-cell patch configuration, mechanically evoked currents were recorded in response to piezo-driven mechanically probing the cell membrane using a blunt glass pipette (Figure 2A). hPIEZO1-transfected cells showed robust probing step-dependent and rapidly inactivating whole-cell inward currents at -80 mV (Figure 2B). The maximal poking current (before the rupture of the cell and loss of the patch) of hPIEZO1 is 995.9±14.0 pA (Figure 2C), and its inactivation Tau is 11.8±1.1 ms (Figure 2D). Remarkably, such mechanically activated currents were recorded neither in vector-transfected nor in hPIEZO1-L322P-transfected HEK-P1-KO cells (Figure 2B-D). These data demonstrate that the L322P mutation completely abolishes mechanically probing-induced whole-cell current.

**Figure 2.**
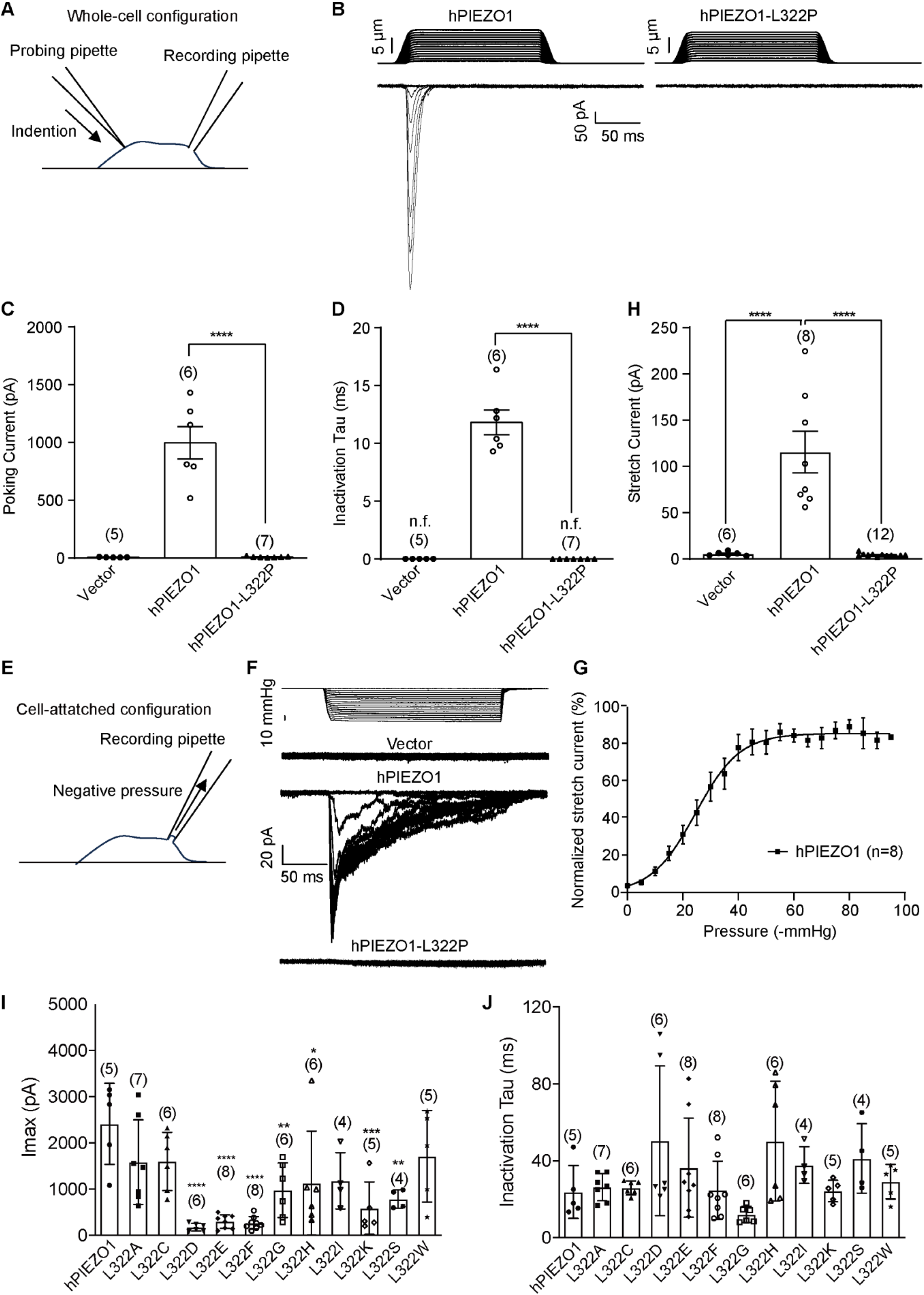
The hPIEZO1-L322P mutation completely abolishes the mechanically activated currents of hPIEZO1. **A**, Schematic representation of the whole-cell patch recording of the mechanically poking-induced currents. **B**, Representative mechanically activated whole-cell current traces from HEK-P1-KO cells heterologously transfected with the indicated constructs in response to the increased poking steps. **C**, Scatter plot of the maximal poking-evoked whole-cell currents mediated by the indicated constructs. The recorded cell numbers are labeled. One-way ANOVA with multiple comparisons. \*\*\*\**p* < 0.0001. **D**, Scatter plot of the inactivation Tau of the poking-evoked whole-cell currents mediated by the indicated constructs. The recorded cell numbers are labeled. One-way ANOVA with multiple comparisons. \*\*\*\**p* < 0.0001. *n.f.* represents no function. **E**, Schematic representation of the cell-attached patch recording of stretch-induced currents. **F**, Representative negative pressure-induced currents from HEK-P1-KO cells transfected with the indicated constructs in response to the increased negative pressure. **G**, The hPIEZO1-mediated pressure-current curve, which is fitted with a Boltzmann equation. **H**, Scatter plot of the maximal stretch-induced current from HEK-P1-KO cells transfected with the indicated constructs. \*\*\*\**p* < 0.0001. **I** and **J**, Scatter plot of the maximal current (**I**) and inactivation Tau (**J**) of the maximal poking-evoked whole-cell current of HEK-P1-KO cells transfected with the indicated constructs. The recorded cell numbers are labeled. One-way ANOVA with Dunn’s comparison to the group transfected with hPIEZO1. \**p* < 0.05, \*\**p* < 0.01, \*\*\**p* < 0.001, \*\*\*\**p* < 0.0001.

To examine whether the hPIEZO1-L322P mutant might respond to other forms of mechanical stimuli, we performed cell-attached patch recording to measure stretch-induced currents by applying negative pressures to the patched cell membrane (Figure 2E). hPIEZO1-transfected cells show negative pressure-dependent increases of the current (Figure 2F-H). Analyzing the pressure-current relationship of hPIEZO1 reveals the half maximal activation pressure (P_50_) of 24.6±7.9 mmHg (Figure 2G) and the maximal stretch-induced current of 115.5 ± 22.4 pA (Figure 2H). By contrast, such stretch-induced currents were recorded neither in vector-transfected nor in hPIEZO1-L322P-transfected HEK-P1-KO cells (Figure 2F, H). Taken together, these results demonstrate that the L322P mutation completely abolishes the mechanical response of hPIEZO1 to membrane stretch.

### The effect of mutating L322 to other amino acids on mechanical activation of hPIEZO1

L322 is located in the middle of TM8 and its change to proline might affect the helical property. To gain more insights into the functional role of L322, we systematically mutated it to other amino acids and characterized their poking-evoked whole-cell currents. We found that mutants including L322D, L322E, L322F, L322G, L322H, L322K and L322S significantly reduced the maximal current amplitude, while other mutants including L322A, L322C, L322I and L322W had no significant effect (Figure 2I). None of the mutations effected the inactivation Tau of the mechanically activated currents (Figure 2J). Together, these data suggest that L322 is required for proper function of hPIEZO1 and its mutation to proline caused the most severe effect in affecting the mechanical activation.

### hPIEZO1-L322P is normally expressed in plasma membrane

PIEZO1 functions as a mechanically activated cation channel in the plasma membrane. The lack of mechanically evoked currents of hPIEZO1-L322P could be due to its lack of proper expression in the plasma membrane. We have previously conducted live immunostaining using the mPIEZO1-FLAG (A2419)-IRES-GFP construct, which has a FLAG-tag inserted after A2419 located in the extracellular cap domain of mPIEZO1 and the IRES-GFP element for reporting positively transfected cells ^23,37^. Following this approach, we inserted the FLAG-tag into the corresponding residue G2399 in the extracellular cap domain of both hPIEZO1 and hPIEZO1-L322P to generate the hPIEZO1-FLAG-IRES-GFP and hPIEZO1-L322P-FLAG-IRES-GFP constructs, respectively. The poking evoked current amplitude and inactivation Tau of hPIEZO1-Flag were similar to those of hPIEZO1 (Figure 3A, B). Furthermore, single-cell Fura-2 calcium imaging revealed their similar responses to the PIEZO1 chemical activator Yoda1 ^38^ (Figure 3C). These data suggest that the FLAG insertion did not alter the channel function of hPIEZO1. Importantly, similar to hPIEZO1-L322P, the hPIEZO1-L322P-FLAG completely lacked mechanically evoked currents (Figure 3A, B). Live immunofluorescent staining without permeabilization of the cell membrane showed that the anti-FLAG immunofluorescent signal in GFP-positive HEK-P1-KO cells transfected with hPIEZO1-FLAG-IRES-GFP was similarly detected as that in hPIEZO1-L322P-FLAG-IRES-GFP-transfected cells (Figure 3D), indicating similar plasma membrane-localized hPIEZO1-FLAG and hPIEZO1-L322P-FLAG proteins. Furthermore, the overall protein level and distribution pattern of hPIEZO1-FLAG and hPIEZO1-L322P-FLAG were similar as revealed by anti-FLAG immunostaining of permeabilized cells (Figure 3E). Under both live and permeabilized staining conditions, no anti-FLAG signal was detected in hPIEZO1-IRES-GFP-transfected cells (Figure 3D, E), demonstrating the specific staining of the FLAG-tag. Taken together, these data demonstrate that the L322P mutation does not affect the expression and proper plasma membrane localization of the hPIEZO1-L322P mutant protein. Thus, the complete loss of mechanically evoked currents of hPIEZO1-L322P is due to its loss of mechanosensitivity.

**Figure 3.**
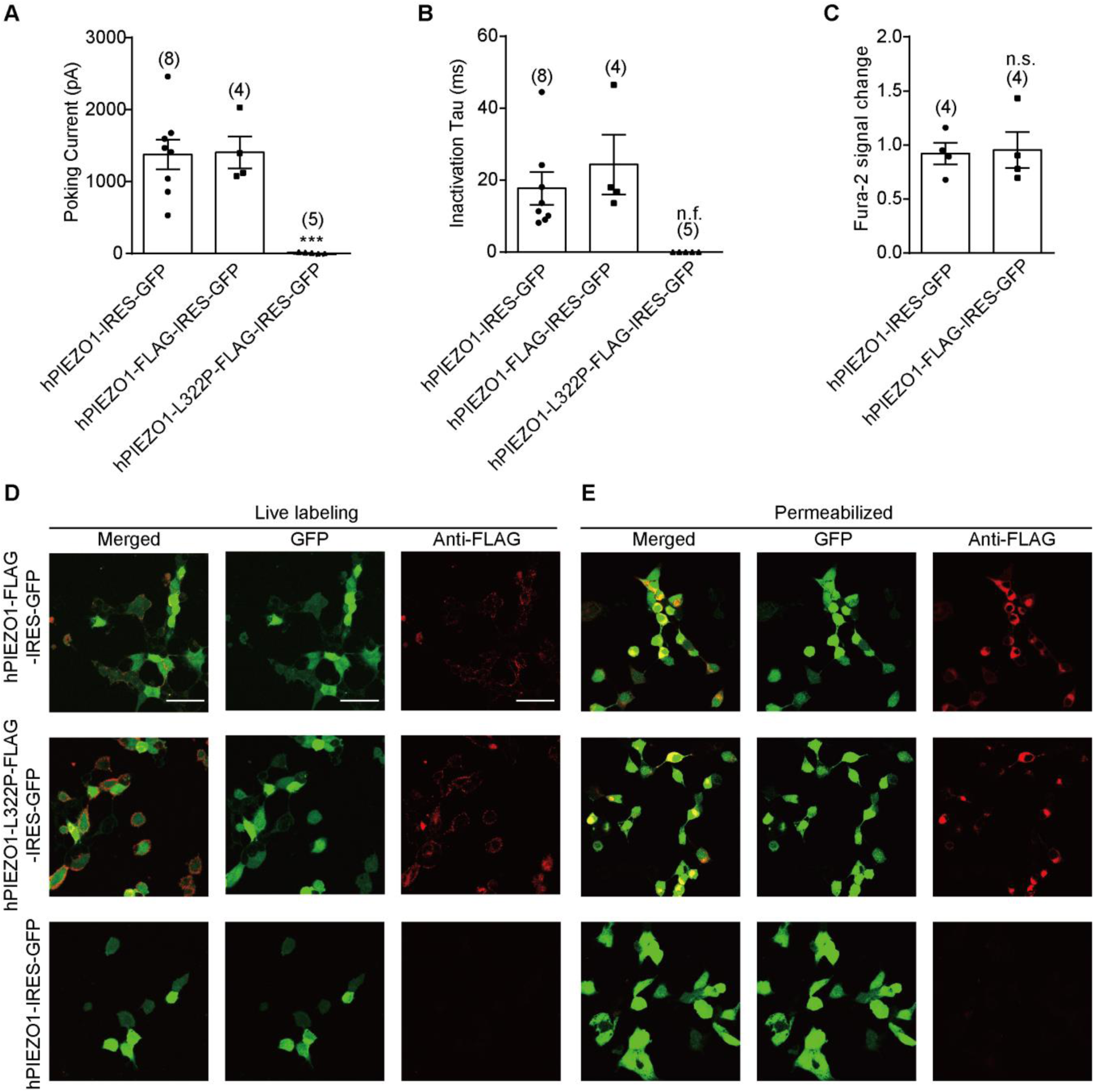
hPIEZO1-L322P is normally expressed in the plasma membrane. **A**, Scatter plot of the maximal step-evoked whole-cell current of HEK-P1-KO cells transfected with the indicated constructs. The recorded cell numbers are labeled. One-way ANOVA with Dunn’s comparison to the group transfected with hPIEZO1-IRES-GFP. \*\*\**p* < 0.001. **B**, Scatter plot of the inactivation Tau of the poking-evoked whole-cell currents of HEK-P1-KO cells transfected with the indicated constructs. The recorded cell numbers are labeled. One-way ANOVA with Dunn’s comparison to the group transfected with hPIEZO1-IRES-GFP. *n.f.* represents no function. **C**, Scatter plot of the Fura-2 ratio change of HEK-P1-KO cells transfected with the indicated constructs in response to the application of 5 μM Yoda1. The recorded coverslip numbers are labeled. Unpaired Student’s test. *n.s.* represents no significance. **D**, Live immunofluorescent staining of HEK-P1-KO cells transfected with the indicated constructs using the anti-FLAG antibody. The extracellularly localized FLAG-tag was inserted after the residue G2399 in the cap domain of human PIEZO1 and hPIEZO1-L322P. The GFP expression under the control of IRES indicates the transfection of the constructs. Scale bar is 50 μm. **E**, Permeabilized immunofluorescent staining of HEK-P1-KO cells transfected with the indicated constructs using the anti-FLAG antibody. Scale bar is 50 μm.

### hPIEZO1-L322P retains normal Ca^2+^ responses to PIEZO1 chemical activators

The proper expression of hPIEZO1-L322P in plasma membrane has prompted us to ask whether it retains the ability to respond to PIEZO1 chemical activators including Yoda1 ^38^ and Jedi1^35^. Using single-cell Ca^2+^ imaging of the Ca^2+^ indicator Fura-2, we found that hPIEZO1 and hPIEZO1-L322P show comparable Fura-2 signal increase in response to the application of either Jedi1 or Yoda1, which was not detected in vector-transfected cells (Figure 4A-C). These data demonstrate that hPIEZO1-L322P retains normal chemical activation.

**Figure 4.**
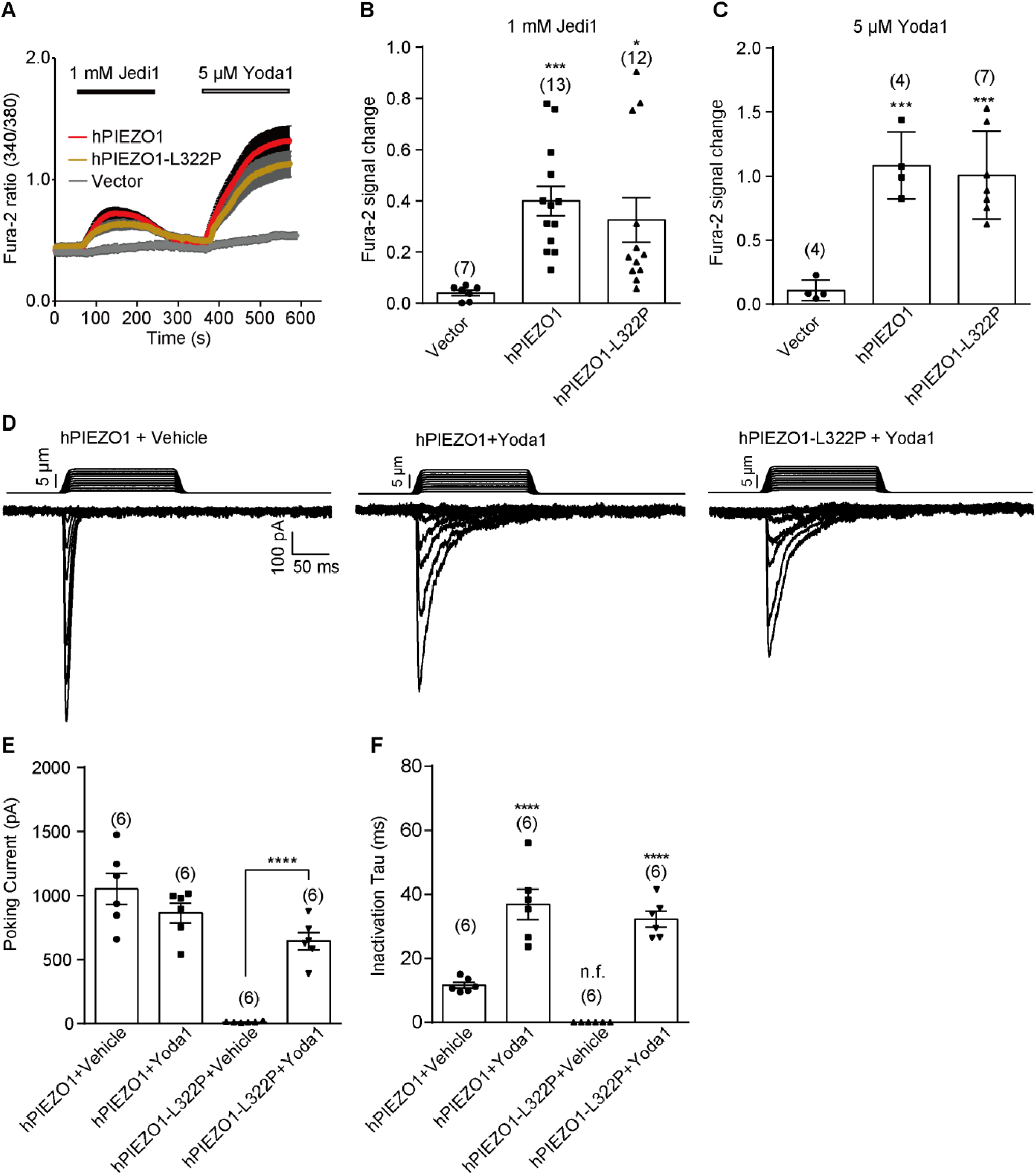
hPIEZO1-L322P shows normal responses to PIEZO1 chemical activators and Yoda1-rescued mechanically activated currents. **A**, Representative averaged single-cell Ca^2+^ imaging traces of HEK-P1-KO cells transfected with the indicated constructs under the indicated treatment conditions. **B**, Scatter plot of the Fura-2 signal ratio change of HEK-P1-KO cells transfected with the indicated constructs in response to the application of 1 mM Jedi1. The recorded numbers of coverslips are labeled. One-way ANOVA with Dunn’s comparison to the group transfected with Vector. \**p* < 0.05, \*\*\**p* < 0.001. **C**, Scatter plot of the Fura-2 signal ratio change of HEK-P1-KO cells transfected with the indicated constructs in response to the application of 5 μM Yoda1. The recorded number of coverslips numbers are labeled. One-way ANOVA with Dunn’s comparison to the group transfected with Vector. \*\*\**p* < 0.001. **D**, Representative poking-evoked whole-cell currents of HEK-P1-KO cells transfected with the indicated constructs in the presence of vehicle or Yoda1. **E**, Scatter plot of the maximal poking step-evoked whole-cell current of HEK-P1-KO cells transfected with the indicated constructs in the presence of vehicle or Yoda1. The recorded cell numbers are labeled. One-way ANOVA with multiple comparisons. \*\*\*\**p* < 0.0001. **F**, Scatter plot of the inactivation Tau of the poking-evoked whole-cell currents of HEK-P1-KO cells transfected with the indicated constructs in the presence of vehicle or Yoda1. The recorded cell numbers are labeled. One-way ANOVA with multiple comparisons. \*\*\**\*p* < 0.0001. *n.f.* represents no function.

### hPIEZO1-L322P shows Yoda1-restored mechanosensitivity

We next tested whether the PIEZO1 chemical activator Yoda1 might restore the mechanosensitivity of hPIEZO1-L322P. Consistent with previously reported effect of Yoda1 on mPIEZO1-mediated poking evoked currents ^35,38^, the presence of Yoda1 slows the inactivation kinetics (Vehicle vs Yoda1: 11.6±0.9 ms vs 36.8±4.7 ms), but does not affect the current amplitude of hPIEZO1 in response to poking stimulation (Figure 4D-F). Remarkably, in the presence of Yoda1, hPIEZO1-L322P showed poking-evoked currents similar to hPIEZO1 (Figure 4D). Both current amplitude (644.6±66.2 pA) and inactivation Tau (32.2±2.4 ms) of hPIEZO1-L322P is comparable to that of hPIEZO1 (Figure 4D-F). By contrast, in the presence of the DMSO vehicle, hPIEZO1-L322P did not show poking evoked currents (Figure 4E, F).

### mPIEZO1-L329P shows similar properties of hPIEZO1-L322P

To examine whether the mutational effect of L322P in human PIEZO1 is conserved in mPIEZO1, we generated the corresponding mutant mPIEZO1-L329P and tested its mechanical and chemical responses. When heterologously expressed in HEK-P1-KO cells, in contrast to mPIEZO1 that mediated both poking- and stretch-induced currents, mPIEZO1-L329P showed no mechanically activated currents (Figure 5A-C). However, mPIEZO1-L329P had similar Yoda1-induced Ca^2+^ response as mPIEZO1 (Figure 5D). These data demonstrate that mPIEZO1-L329P had abolished mechanical responses, but retained Yoda1-evoked chemical response, which is reminiscent of hPIEZO1-L322P.

**Figure 5.**
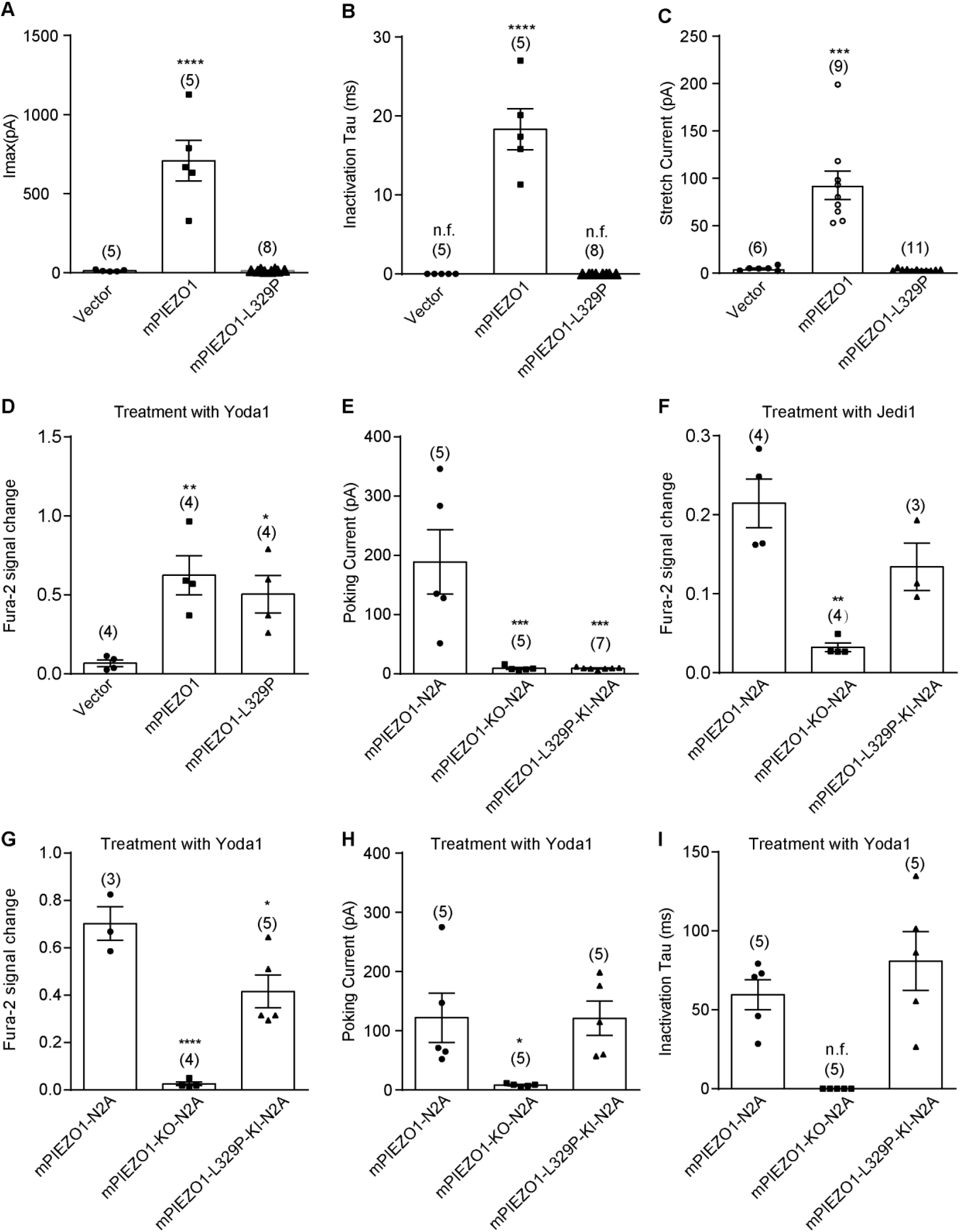
Heterologously expressed or endogenously knocked-in mPIEZO1-L329P specifically lacks mechanical response, but retains normal responses to PIEZO1 chemical activators and Yoda1-rescued mechanically activated currents. **A**, Scatterplot of the maximal poking-evoked whole-cell currents from HEK-P1-KO cells transfected with the indicated constructs. The recorded cell numbers are labeled. One-way ANOVA with Dunn’s comparison to the group transfected with vector. \*\*\*\**p* < 0.0001. **B**, Scatter plot of the inactivation Tau of the poking-evoked whole-cell currents from HEK-P1-KO cells transfected with the indicated constructs. The recorded cell numbers are labeled. One-way ANOVA with Dunn’s comparison to the group transfected with vector. \*\*\*\**p* < 0.0001. *n.f.* represents no function. **C**, Scatterplot of the stretch-induced currents from HEK-P1-KO cells transfected with the indicated constructs. The recorded cell numbers are labeled. One-way ANOVA with Dunn’s comparison to the group transfected with vector. \*\*\**p* < 0.001. **D**, Scatter plot of the Fura-2 signal ratio change of HEK-P1-KO cells transfected with the indicated constructs in response to the application of 5 μM Yoda1. The recorded numbers of coverslips are labeled. One-way ANOVA with Dunn’s comparison to the group transfected with Vector. \**p* < 0.05, \*\**p* < 0.01. **E**, Scatterplot of the maximal poking-evoked whole-cell currents from wild-type N2A cells (mPIEZO1-N2A), N2A cells with the endogenous PIEZO knocked out (mPIEZO1-KO-N2A), or N2A cells with the L329P mutation knocked into the endogenous PIEZO1 (mPIEZO1-L329P-KI-N2A). The recorded cell numbers are labeled. One-way ANOVA with Dunn’s comparison to the mPIEZO1-N2A group. \*\*\**p* < 0.001. **F**, Scatter plot of the Fura-2 signal ratio change of mPIEZO1-N2A, mPIEZO1-KO-N2A, mPIEZO1-L329P-KI-N2A cells in response to the application of 1 mM Jedi1. The recorded number of coverslips numbers are labeled. One-way ANOVA with Dunn’s comparison to the mPIEZO1-N2A group. \*\**p* < 0.01. **G**, Scatter plot of the Fura-2 siganl ratio change of mPIEZO1-N2A, mPIEZO1-KO-N2A, mPIEZO1-L329P-KI-N2A cells in response to the application of 30 μM Yoda1. The recorded number of coverslips numbers are labeled. One-way ANOVA with Dunn’s comparison to the mPIEZO1-N2A group. \**p* < 0.05, \*\*\*\**p* < 0.0001. **H**, Scatter plot of the maximal poking step-evoked whole-cell current of mPIEZO1-N2A, mPIEZO1-KO-N2A, mPIEZO1-L329P-KI-N2A cells in the presence of 30 μM Yoda1. The recorded cell numbers are labeled. One-way ANOVA with Dunn’s comparison to the mPIEZO1-N2A group. \**p* < 0.05. **I**, Scatter plot of the inactivation Tau of the poking-evoked whole-cell currents of mPIEZO1-N2A, mPIEZO1-KO-N2A, mPIEZO1-L329P-KI-N2A cells in the presence of 30 μM Yoda1. The recorded cell numbers are labeled. One-way ANOVA with Dunn’s comparison to the mPIEZO1-N2A group. *n.f.* represents no function.

### Endogenous mPIEZO1-L329P mutant shows defective but Yoda1-rescuable mechanical responses

We further examined the mutational effect of L329P in endogenously expressed PIEZO1 in the mouse neuron-2A (N2A) cell line, in which mPIEZO1 was originally cloned^1^. Toward this, we used the CRISPR/Cas9 technology to either knock out the PIEZO1 gene or knock in the L329P mutation into the *PIEZO1* gene in N2A cells, generating the mPIEZO1-KO-N2A (Supplementary Figure 2A, B) and PIEZO1-L329P-KI-N2A (Supplementary Figure 2C-E) cell lines, respectively. We detected poking-induced currents and Jedi1- or Yoda1-induced Ca^2+^ responses in N2A cells, which were absent in the mPIEZO1-KO-N2A cells, demonstrating endogenous mPIEZO1-mediated mechanical and chemical responses in N2A cells (Figure 5E, F, G). Importantly, the PIEZO1-L329P-KI-N2A cells lacked poking evoked currents, but showed Jedi1- or Yoda1-induced Ca^2+^ responses (Figure 5E, F, G). Remarkably, Yoda1 restored the mechanically activated currents in the mPIEZO1-L329P-KI-N2A cells with an averaged current of 53.5±1.5 pA and an inactivation Tau of 72.9±13.6 ms, which were similar to the current recorded in N2A cells (42.2±4.7 pA and 41.5±8.4 ms) (Figure 5H, I). By contrast, no mechanically activated currents were observed in the PIEZO1-KO-N2A cells even in the presence of Yoda1 (Figure 5H, I). These data demonstrate that the endogenously expressed mPIEZO1-L329P mutant lacks direct responses to mechanical stimulation, but retains the ability to respond to PIEZO1 chemical activators and gain Yoda1-restored mechanosensitivity.

## Discussion

Here we discover that the inherited compound heterozygous variants of *PIEZO1* :c.2991+1 G>A; p.E997 and c.965 T>C; p. L322P led to recurrent fetal hydrops and abortion (Figure 1A). The splicing site variant c.2991+1 G>A is predicted to generate the hPIEZO1-E997-truncation protein lacking the C-terminal pore domain, and thus is expected to cause LOF of PIEZO1 channel. The missense variant c.965 T>C generates the single-residue mutation hPIEZO1-L322P. On the basis of a combination of electrophysiological, Ca^2+^ imaging and immunostaining characterizations, we find that the hPIEZO1-L322P mutant completely lacks mechanically evoked currents, but shows normal expression and localization in the plasma membrane and responses to the PIEZO1 chemical activators Yoda1 and Jedi1 (Figures 2-4). Thus, our data demonstrate that the L322P mutation specifically abolishes the mechanosensitivity of PIEZO1, but does not affect its proper expression and organization into functional channels with normal responses to chemical activations. Previous studies have linked LOF of PIEZO1 due to either nonsense or splicing site mutations to autosomal recessive GLD with non-immune hydrops fetalis ^19^. Taken together, we conclude that hPIEZO1-L322P represents a specific loss-of-mechanosensitivity mutant of PIEZO1 that directly contributes to the pathological phenotype of fetal hydrops, providing compelling evidence supporting the critical role of the mechanosensitivity of PIEZO1 in determining its in vivo pathophysiological functions involving mechanotransduction. Given that similar changes of properties have been consistently observed in the mPIEZO1-L329P mutant either heterologously overexpressed or endogenously knocked into N2A cells (Figures 5), it will be interesting to generate the mPIEZO1-L329P knock-in mouse model for dissecting out the specific role of the mechanosensitivity of mPIEZO1 in determining its diverse physiological and pathophysiological functions.

Mechanistically, it is remarkable that the single residue mutation L322P or L329P in the large hPIEZO1 or mPIEZO1 of 2522 or 2547 residues, respectively, completely abolishes the mechanosensitivity of PIEZO1. We have previously found that deletion of the extracellular loop connecting TM15-16 (EL15-16) of the THU4 or EL19-20 of the THU5 greatly reduced the mechanically poking or stretching induced currents of mPIEZO1 ^23^. Intriguingly, these deletion mutants of mPIEZO1 also abolish the response of the hydrophilic and extracellular acting Jedi1 and Jedi2, but retain normal responses to the hydrophobic Yoda1^35^. These studies have led to the proposal that the blade domain together with the intracellular long beam domain functions as a lever-like apparatus as a designated transduction pathway for long-distance mechano- and chemical-gating of the downstream C-terminal pore^35^. L322 is located in the distal TM8 of the THU2 of the blade domain, further upstream of THU4 and THU5. The complete lack of mechanosensitivity of hPIEZO1-L322P and mPIEZO1-L329P is consistent with the assigned mechanosensing function of the blade domain. THU1-3 of PIEZO1 is highly flexible and its lateral expansion has been associated with mPIEZO1 activation ^23,25,26,39^. We speculate that the L322P mutation might affect the ability of the distal THU1-3 for mechanical activation of PIEZO1. Interestingly, hPIEZO1-L322P and mPIEZO1-L329P retain the ability to normally respond to both Jedi1 and Yoda1. These data suggest that Jedi1 and Yoda1 function at downstream of the L322-residing mechanosensing sub-domain, supporting the mechanotransduction pathway from the most distal part of the blade domain to the central pore.

It is striking that the complete loss of mechanosensitivity of hPIEZO1-L322P and mPIEZO1-L329P can be effectively rescued by Yoda1. Yoda1 and Jedi1/2 have been shown to potentiate the mechanosensitivity of PIEZO1 ^35,38^, which might explain their ability to rescue the mechanosensitivity of hPIEZO1-L322P and mPIEZO1-L329P. Notably, mPIEZO1 mutants lacking EL15-16 or EL19-20 lost response to mechanical stimulation and Jedi1/2 chemical activation, but have normal responses to Yoda1 and Yoda1-rescuable mechanical responses ^23^. These findings suggest that Yoda1 might function at the downstream of the L322-EL15-16-EL19-20-beam-pore mechanotranduction pathway, in which Jedi1/2 might function in between L322 and EL15-16. In line with this, Yoda1 has been proposed to bind to a pocket in between THU8 and THU9, which is close to the pore ^40,41^. Notably, literally reported GLD-associated missense mutations of hPIEZO1 are scattered from the blade to the pore ^7,21^. Thus, the revelation of the full-chain mechanotransduction pathway and functional characterizations of the loss-of-mechanosensitivity and disease-causing mutations might allow the use of different PIEZO1 chemical activators such as Jedi1/2 and Yoda1 for restoring their mechanosensitivity, offering therapeutic potential for precise treatment of human patients with the associated genetic diseases such as GLD with fetal hydrops.

## Methods

### Molecular genetic analysis

The human genetic study was approved by the institutional ethics committee. The parents agreed to participate in the study and provided signed informed consent. For the Chromosome Number Variation sequencing (CNVseq), genomic DNA was extracted from fetus skin tissue using QIAamp DNA Mini Kit following the manufacturer’s protocol. The DNA was fragmented to 300-500 bp using the Genomic DNA Fragmentation Kit from Jiabao Renhe Company. Adaptors for PCR and Illumina sequencing were ligated by T4 DNA ligase, followed by PCR amplification with specific primers and cycling conditions. The library was purified with Agencourt AMPure XP beads. Sequencing was performed on the Illumina NextSeqTM Dx550 with 150bp paired - end reads and a 1-5X mean depth. Raw reads were quality - controlled, with low- quality ones and adaptor sequences removed. High - quality reads were aligned to the human reference genome (hg19 or hg38) using BWA. Read depth was calculated by BEDTools, and copy number changes were detected by comparing with normal control samples. CNVnator was used to identify potential CNVs, which were further annotated with the Database of Genomic Variants (DGV) to determine novelty and functional implications.

For the Trio-based whole-exome sequencing , the DNA library was constructed using a Roche SeqCap EZ MedExome Enrichment kit, and high-throughput sequencing was conducted on an Illumina HiSeq X platform, raw FASTQ data were generated and processed through the subsequent steps: a) map the short reads to a reference genome (hg19) using the BWA software (v0.7.12-r1039) ^42^; b) use the SAMtools software (v0.1.18) to sort the short sequences and convert the format of the data; c) use the Picard software (v1.134) (http://broadinstitute.github.io/picard/) to mark duplicate reads; d) use the Genome Analysis Toolkit (GATK v3.7) ^43^ to identify SNVs and indels; e) perform functional annotation of these variant sites using the ANNOVAR software (16 Jul 2016 version). All identified sequence variants were systematically assessed and classified based on the standards and guidelines set forth by the American College of Medical Genetics and Genomics (ACMG)^44^.

For variants PCR amplification and Sanger Sequencing, Genomic DNA were subjected to variants PCR amplification and Sanger sequencing. The nucleotide sequence of the DNA fragment containing the mutation site was retrieved from the UCSC Genome Browser. Using Primer 3.0, a specific primer was designed to target this region for amplification. The corresponding gene fragment was amplified from the genomic DNA samples using polymerase chain reaction (PCR). The amplified DNA fragments were subsequently verified through Sanger sequencing. Nucleotide variations in the aligned sequencing reads were identified and reviewed using NextGENe software (SoftGenetics, State College, PA).

### Molecular cloning

The human PIEZO1 (hPIEZO1) and mouse PIEZO1 (mPIEZO1) were kindly provided by Dr. Ardem Patapoutian at Scripps Research Institute. The WT and mutated PIEZO1 cDNAs were cloned into the pcDNA3.1 expression vector. To indicate the expression of target protein, the encoding sequences for wild-type hPIEZO1, hPIEZO1-L322P were followed by the GFP-coding sequence and driven by an internal ribosome entry site (IRES) to monitor transfection efficiency and protein expression. The WT mPIEZO1 and mPIEZO1-L329P mutants were directly fused to the mRuby2 fluorescent protein-encoding sequence to accurately reflect the expression and subcellular localization of the proteins. All the PIEZO1-related constructs including hPIEZO1-L322P-IRES-GFP, mPIEZO1-L329P-mRuby2, hPIEZO1-FLAG-IRES-GFP, hPIEZO1-L322P-FLAG-IRES-GFP and other molecular cloning were subcloned by using the One Step Cloning Kit according to the instruction manual (Vazyme Biotech) as previous described^45,46^. All constructs were validated through sequencing.

### Cell culture

All the electrophysiology tests were performed in the human embryonic kidney 293T in which the endogenous PIEZO1 gene was disrupted (HEK-P1-KO cells) (gift from Dr. Ardem Patapoutian Laboratory). The HEK-P1-KO cells were cultured in DMEM supplemented with 10% FBS and 1% P/S, and transfected with Lipofectamine 3000 (Invitrogen).

### Immunostaining

For live-cell labelling, cells cultured on coverslips were treated with the anti-Flag antibody (1:100, Sigma) mixed in prewarmed culture medium and incubated at room temperature for 1 h. Following three washes, the cells were treated with the Alexa Fluor 594 donkey-anti-mouse IgG secondary antibody (1:200, Life Technologies) at room temperature for 1 h, washed again, and subsequently fixed with 4% (w/v) paraformaldehyde. For permeabilized staining, the cells were initially fixed with 4% (w/v) paraformaldehyde, permeabilized using 0.2% (w/v) Triton X-100, and then incubated with the anti-Flag antibody (1:200, Sigma) at room temperature for 1 h. The cells were washed and subsequently incubated with the Alexa Fluor 594 donkey-anti-mouse IgG (1:200, Life Technologies) or Alexa Fluor 594 donkey-anti-rabbit IgG (1:200, Life Technologies) secondary antibody at room temperature for 1 h. After another round of washing, the coverslips were mounted and imaged under a Nikon A1 confocal microscope equipped with a 60× oil objective (N.A. 5 1.49), using excitation wavelengths of either the 488 nm (GFP channel) or 561 nm (TRITC channel).

### Generation of the mPIEZO1-KO-N2A and mPIEZO1-L329P-KI-N2A cell lines

To generate the mPIEZO1-KO-N2A cell lines, we targeted the intronic regions flanking exon 39 of mPIEZO1 (upstream and downstream introns) to precise excise exon 39, resulting in a frameshift mutation due to the removal of non-integer multiple of 3 nucleotides. The KO cell line was validated by genomic PCR and Sanger sequencing.

To generate the mPIEZO1-L329P-KI-N2A cell lines, the single-guide RNA (sgRNA) targeting the L329 in exon 8 of PIEZO1 gene locus was designed by using the CRISPR Design Tool (http://crispr.mit.edu/). The sgRNA oligos were cloned into the pX458-GFP backbone via Golden Gate Assembly. Transformation was performed using XL10-gold chemically competent cell, followed by a 24 h incubation. Plasmid DNA was purified using Mini Plasmid Kit (TIANGEN, DP103) and analyzed by Sanger sequencing. The donor DNA for knock-in was synthesized by homologous recombination. PCR synthesis was carried out by using mouse genomic DNA and pCDNA3.1(-)-mPiezo1-mRuby as the amplification templates. The purified sgRNA and donor DNA were mixed together and transfected into the cultured Neuro-2A cell dish. Single cell clones were isolated and the genotypes of cell lines were identified by genomic PCR following Sanger sequencing. The sgRNA sequence, donor DNA sequence, primers for genomic PCR and Sanger sequencing were listed in the Supplementary Table 1.

### Whole-cell Electrophysiology

All the electrophysiology tests were performed on HEK-P1-KO cells in which the endogenous PIEZO1 gene is deleted. Cells were transfected with constructs PIEZO1 or variant fused with GFP or mRuby using transfection reagent Lipofectamine 3000, and the GFP or mRuby fluorescence signals were assessed 24 h post-transfection to evaluate transfection and expression efficiency. Cells were gently digested with a diluted 0.05% trypsin solution (1:20 ratio), sparsely replated onto poly-D-lysine-coated coverslips, and incubated approximately 2 h before recording. Patch-clamp experiments were performed using the HEKA EPC10 system, following previously established protocols ^22,23^. For whole-cell patch-clamp recordings, the recording electrodes, filled with an internal solution containing 133 mM CsCl, 1 mM CaCl_2_, 1 mM MgCl_2_, 5 mM EGTA, 10 mM HEPES (adjusted to pH 7.3 with CsOH), 4 mM MgATP and 0.4 mM Na_2_GTP, exhibited a resistance ranging from 2 to 5 MΩ. The extracellular solution consisted of 133 mM NaCl, 3 mM KCl, 2.5 mM CaCl_2_, 1 mM MgCl_2_, 10 mM HEPES (adjusted to pH 7.3 with NaOH) and 10 mM glucose. All experiments were performed at room temperature. Currents were sampled at a rate of 20 kHz and filtered at 2 kHz using Patchmaster software. Leak currents recorded prior to mechanical stimulation were subtracted offline from the current traces. Mechanical stimulation was applied to the cell during patch-clamp recording at an 80° angle using a fire-polished glass pipette with a tip diameter of 3–4 μm, as described in previous protocols ^1^. The downward movement of the probe towards the cell was controlled by a Clampex-operated piezoelectric crystal micro-stage (E625 LVPZT Controller/Amplifier; Physik Instrument). The probe moved at a velocity of 1 μm.ms^−1^ during both downward and upward motions, with the stimulus duration set at 150 ms. Mechanical steps were applied in 1 μm increments at intervals of 8 s, and currents were recorded at a holding potential of −80 mV.

### Cell-attached electrophysiology

Stretch-activated currents were measured using the standard cell-attached patch-clamp configuration with the HEKA EPC10 system, following established protocols^2,45^. Currents were sampled at 20 kHz and filtered at 2 kHz. The recording electrodes, filled with a standard solution containing (in mM) 130 NaCl, 5 KCl, 10 HEPES, 1 CaCl2, 1 MgCl2 and 10 TEA-Cl (adjusted to pH 7.3 with NaOH), had a resistance of 2-3 MΩ. The external solution, used to zero the membrane potential, was composed of (in mM) 140 KCl, 10 HEPES, 1 MgCl2, and 10 glucose (adjusted to pH 7.3 with KOH). All experiments were performed at room temperature. Membrane patches were stimulated with 500 ms negative-pressure pulses delivered through the recording electrode using a Patchmaster-controlled pressure clamp HSPC-1 device (ALA-Scientific). Stretch-activated channels were recorded at a holding potential of 120 mV, with pressure steps ranging from 0 to 100 mm Hg in increment of 10 mm Hg. For each cell, 4 to 11 recording traces were averaged for analysis. The current-pressure relationships were fitted using a Boltzmann equation of the form: *I(P)* = [1 + exp (-(*P* – *P_50_*)/*s*)]^− 1^, where *I* represents the peak stretch-activated current at a given pressure, *P* is the applied patch pressure (in mm Hg), *P_50_* is the pressure required for half-maximal channel activation, and *s* reflects the current’s sensitivity to pressure.

### Single-cell Fura-2 Ca^2+^ Imaging

HEK293T cells cultured on 8-mm round glass coverslips coated with poly-D-lysine and positioned in 48-well plates were transfected with either hPIEZO1 or hPIEZO-L322P. Subsequently, these cells were subjected to Fura-2 Ca^2+^imaging 36h post transfection, following established protocols^35^. The confluent cells were rinsed with a buffer containing 1 × HBSS (supplemented with 1.3 mM Ca^2+^) and 10 mM HEPES (adjusted to pH 7.2 with NaOH), then incubated at room temperature for 30 min with 2.5 μM Fura-2-AM (Molecular Probes) and 0.05% Pluronic F-127 (Life Technologies). Following the buffer wash, the coverslips were positioned onto an inverted Nikon-Tie microscope equipped with a CoolSNAP charge-coupled device (CCD) camera and Lambda XL light box (Sutter Instrument). GFP -positive and -negative cells were chosen for the measurement of the 340/380 ratio using a 20× objective (numerical aperture N.A. = 0.75) and the MetaFluor Fluorescence Ratio Imaging software (Molecular Device). A stock solution of Yoda1 at 30 mM was dissolved in DMSO and subsequently diluted to a final concentration of 5 μM. The compound solutions were delivered into the chamber and to the cells via a multichannel perfusion system (MPS-2, World Precision Instruments).

### Data Analysis and Statistics

All data are shown as mean±standard error of mean (S.E.M). Data normality was tested using the D’Agostino-Pearson omnibus normality test or the Shapiro-Wilk normality test or Kolmogorov-Smirnov test. For data that followed a normal distribution, statistical significance was assessed using either an unpaired Student’s t-test for comparisons between two groups, or one-way ANOVA for comparisons involving multiple groups. Statistical significance were indicated as: \**p* < 0.05, \*\**p* < 0.01, \*\*\**p* < 0.01, \*\*\*\**p* < 0.0001.

## Acknowledgments

We thank Dr. Yaxiong Cui and Gewei Zhou for help and discussion. This work was supported by grant numbers 2021ZD0203301, 32130049, 32425003, 32021002, 31825014 from either the National Key R&D Program of China or the National Natural Science Foundation of China, and the New Cornerstone Investigator Program, Beijing Outstanding Young Scientist Program grant, Research Fund of Vanke School of Public Health and Tsinghua University Initiative Scientific Research Program to B.X.

## Author Contributions

J.J. performed molecular cloning, biochemical characterizations, electrophysiology, calcium imaging and data analysis; W.G. carried out case collection and sequencing; X.D. did biochemical characterizations; Y.W. generated the mPIEZO1-KO-N2A cell line; J.Q., L.Y. and X.Z. supervised clinical studies; B.X. conceived and directed the study, wrote the manuscript with J.J.; All authors read, discussed and edited the manuscript.

## Competing interests

The authors declare no competing interests.

## Author Information

Correspondence and requests for materials should be addressed to B.X. and J.Q.

## Supplementary Figures and Legends

**Supplementary Figure 1.**
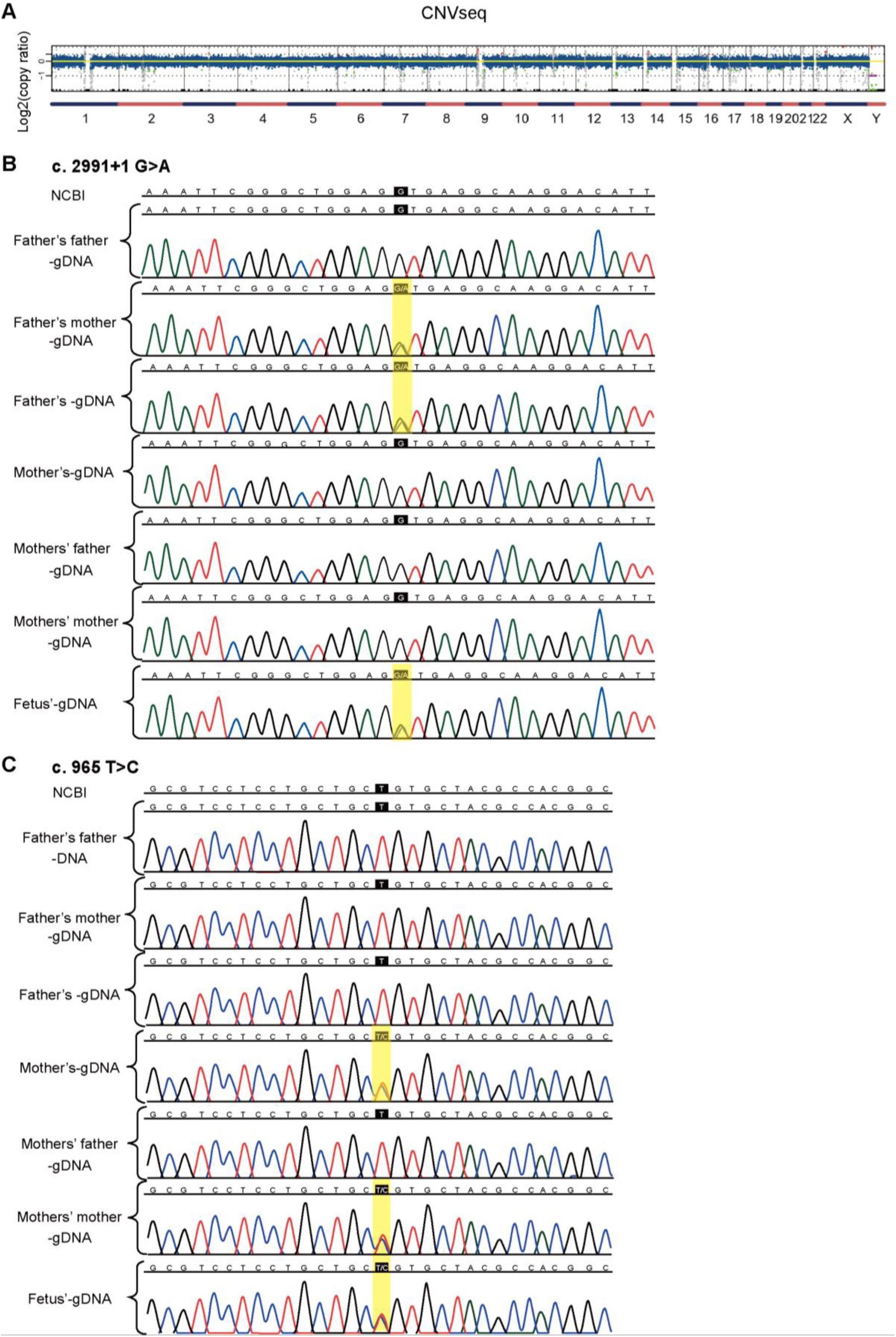
Identification of the fetal hydrops-associated mutations of PIEZO1. **A**, CNV-seq analysis of the aborted fetus. **B**, Alignment of exome sequencing reads from all individuals to the reference human genome illustrates the c.2991+1 G>A mutation of *hPIEZO1*. Variant bases are highlighted in yellow in the sequencing reads. The reference sequence is indicated at the top of the imaging. **C**, Alignment of exome sequencing reads from all individuals to the reference human genome illustrates the c.965 T>C mutation. Variant bases are highlighted in yellow in the sequencing reads. The reference sequence is indicated at the top.

**Supplementary Figure 2.**
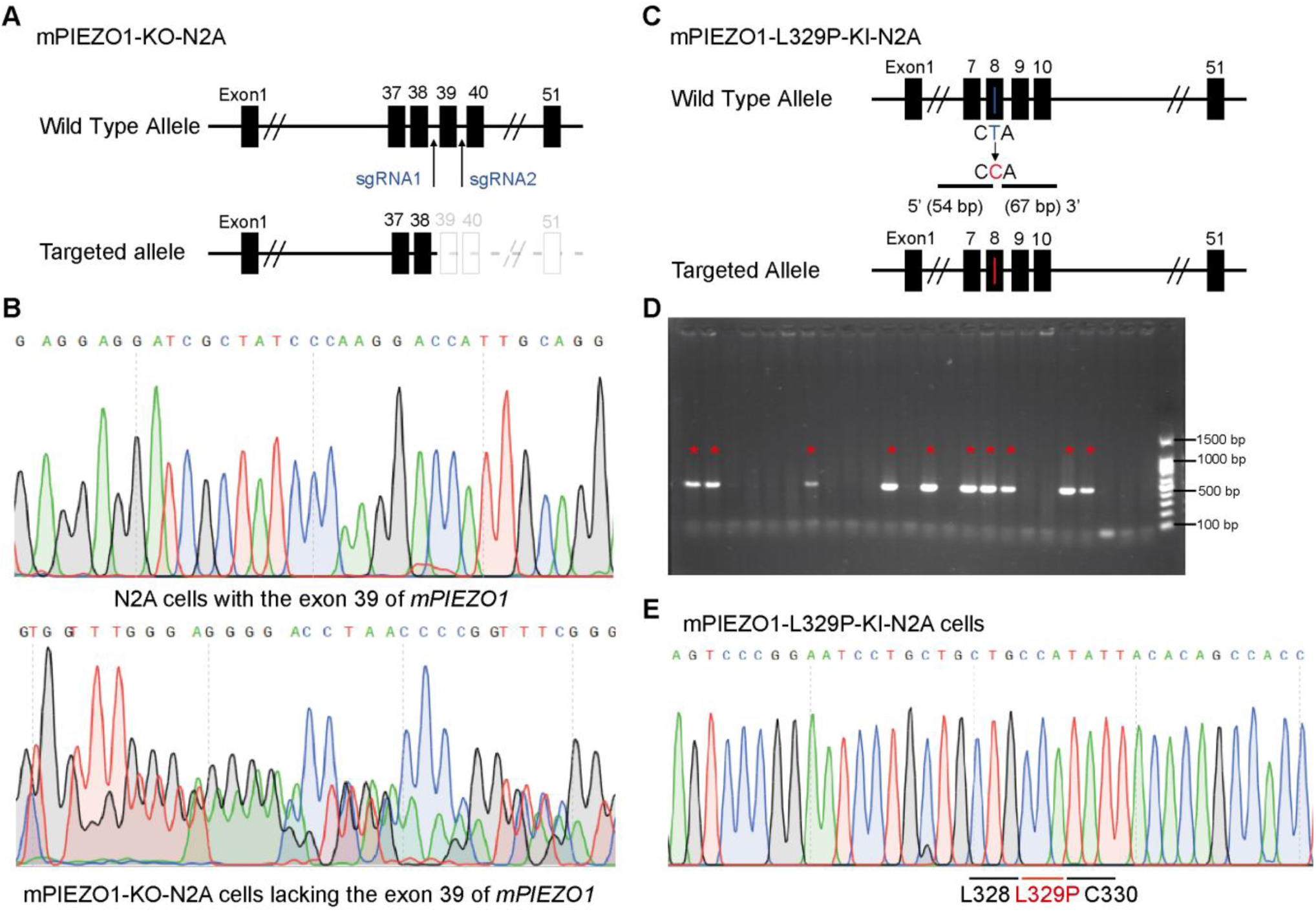
Generation and characterization of the mPIEZO1-KO or mPIEZO1-L329P-knock-in N2A cells. **A**, Schematic representation of the CRISPR-Cas9-mediated deletion of the endogenous *mPIEZO1* gene in N2A cells. **B**, DNA sequencing verification of the DNA sequence corresponding to the exon 39 of mPIEZO1 gene in wild type N2A cells or mPIEZO1-KO-N2A cells. **C**, Schematic representation of the CRISPR-Cas9-mediated knock-in of the L329P mutation (by changing the codon from CTA to CCA) into the endogenous *mPIEZO1* gene in N2A cells. **D**, PCR screening of mPIEZO1-L329P-KI-N2A clones after the CRISPR-Cas9-mediated genetic targeting. Clones labeled with * show the expected PCR product. **E**, DNA sequencing verification of the correct insertion of the L329P-encoding sequence (mutation from CTA to CCA).

**Supplementary Table 1.**
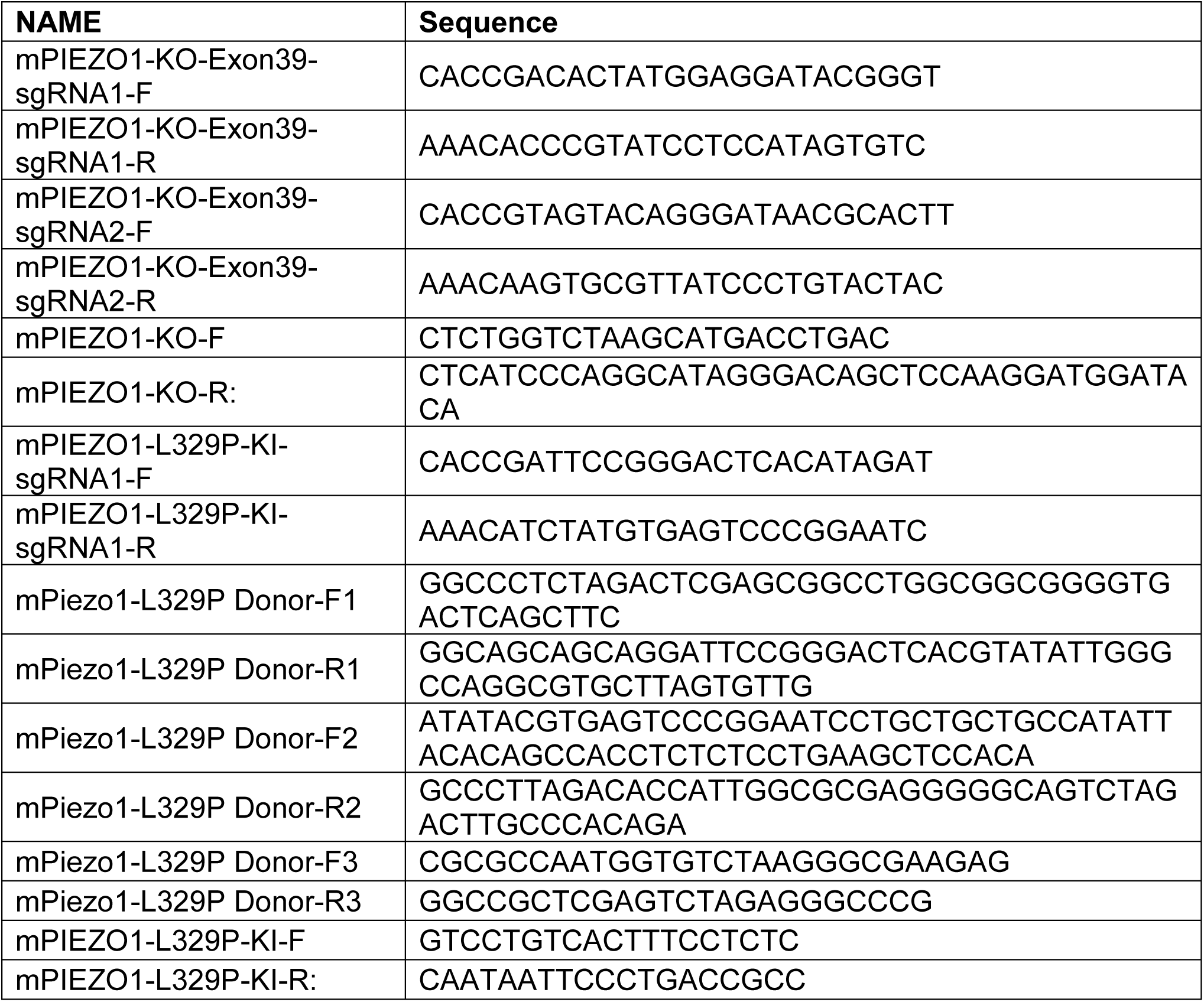
Primers and sgRNA sequences.

## Notes

### Competing Interest Statement

The authors have declared no competing interest.

